# Automated detection of sleep-boundary times using wrist-worn accelerometry

**DOI:** 10.1101/225516

**Authors:** Johanna O’Donnell, Sven Hollowell, Gholamreza Salimi-Khorshidi, Carmelo Velardo, Claire Sexton, Kazem Rahimi, Heidi Johansen-Berg, Lionel Tarassenko, Aiden Doherty

**Affiliations:** Institute of Biomedical Engineering, Department of Engineering Science, University of Oxford, Oxford, United Kingdom; George Institute for Global Health, Nuffield Department of Obstetrics and Gynaecology, University of Oxford, Oxford, United Kingdom; Nuffield Department of Population Health, University of Oxford, Oxford, United Kingdom; Nuffield Department of Neuroscience, University of Oxford, Oxford, United Kingdom

**Author notes:** These authors are joint last authors (Lionel Tarassenko, Aiden Doherty). Current Address: Nuffield Department of Population Health, Richard Doll Building, Old Road Campus, Headington, Oxford OX3 7LF, United Kingdom.

## Abstract

**Objective:** Current polysomnography-validated measures of sleep status from wrist-worn accelerometers cannot be used in fully automated analysis as they rely on self-reported sleep-onset and -end (sleep-boundary) information. We set out to develop an automated, data-driven approach to sleep-boundary detection from wrist-worn accelerometer data.

**Methods:** On three separate occasions, participants were asked to wear a GENEActiv® wrist-worn accelerometer for nine days and concurrently complete sleep diaries with lights-off, asleep and wake-up information. We developed and evaluated three data-driven methods for sleep-boundary detection: a change-point detection based method, a thresholding method and a random forest classifier based method. Mean absolute errors between automatically-derived and self-reported sleep-onset and wake-up times were recorded in addition to kappa statistics for the minute-by-minute performance of each of the methods.

**Results:** 46 participants provided 972 days of accelerometer recordings with corresponding self-reported sleep information. The three sleep-boundary detection methods resulted in mean absolute errors in sleep-onset and wake-up times per individual of 36 min, 34 min and 33 min and kappa statistics of 0.87, 0.89 and 0.89, respectively.

**Conclusion:** Our methods provide a data-driven approach to detect sleep-onset and -end times without the need for self-reported sleep-boundary information. The methods are likely to be of particular use for large-scale studies where the collection of self-reported sleep diaries is impractical.

**Significance:** Objective measures of sleep are needed to reliably detect associations with health outcomes. This work lays the foundation for studies of objectively measured sleep duration and its health consequences in large studies.

## Introduction

Alterations in sleep duration and changes in sleep-wake timing are associated with a wide range of negative health outcomes, including an increased risk of type 2 diabetes and cardiovascular outcomes [1–3] as well as increased incidence of psychiatric disorders [4]. However, evidence often relies on self-reported sleep information, which may be unreliable and affected by measurement errors due to memory bias [5]. As a result, many studies now use wrist-worn accelerometers in an attempt to objectively measure sleep durations [6, 7]. Publicly available sleep detection methods developed for wrist-worn accelerometers include the idleness-detecting method by *Borazio et al* [8] and the angular-movement based method by *van Hees et al* [6]. Methods such as these have traditionally been developed in a laboratory environment and tested against the gold standard for objective sleep analysis, polysomnography.

However, current polysomnography-validated measures of sleep status from wrist-worn accelerometers cannot be used in fully automated analysis as they rely on self-reported sleep-onset and –end (sleep-boundary) information [6]. This information is typically collected through means of a sleep diary [6] or through a button on the accelerometer which participants can press as they go to bed and wake up [5]. As a result it is not feasible to collected self-reported sleep-boundary times for all participants in large scale studies, such as the UK Biobank [7], and therefore a need exists for an automated way to extract sleep-onset and wake up times.

In this paper we set out to develop a fully-automated method for sleep-boundary detection from wrist-worn accelerometer data. We also set out to assess its performance in free-living scenarios in healthy UK adults aged 60-80.

## Materials and methods

### Study Population

Accelerometer recordings and self-reported sleep-boundary data were collected as part of a Cognitive Health in Ageing (CHA) Exercise study [9]. The purpose of this study was to analyse the effect of an anaerobic exercise intervention on brain MRI measures. Adults between the ages of 60 and 80 years that self-reported less than 60 minutes of heart-raising physical activity per week and no contraindications to MRI scanning or fitness testing were eligible to take part in the research study. Ethical Approval for this study was obtained from the Local Research Ethics Committee (Oxford REC B Ref 10/H0605/48).

### Study Procedure

Participants wore the GENEActiv® accelerometer on their non-dominant wrist for nine consecutive days in weeks 0, 12 and 24 of the study and filled in daily sleep diaries with lights-off, asleep and wake-up information during these periods. The GENEActiv® sensor contains a tri-axial accelerometer, with a sensor resolution of +-8g, as well as a temperature sensor and has previously been validated for sleep research [10] [6]. Acceleration data were collected at a sampling frequency of 87.5 Hz.

### Data Preparation

Accelerometer recordings with at least one day of corresponding asleep and wake-up labels were selected for the analysis. Accelerometer data followed careful UK Biobank preprocessing and quality control checks [7], including device calibration to local gravity [11], removal of machine noise and gravity [12], and imputation of non-wear data segments using the average of similar time-of-day vector magnitude data points from different days of the measurement. Device non-wear time was automatically identified as consecutive stationary episodes lasting for at least 60 minutes [7].

### Sleep-boundary Detection

In this study, we employ three approaches for automatic sleep-boundary detection: (1) A statistical technique for detecting change points in the accelerometer time series, (2) A data-driven thresholding method to classify short intervals into sleep and wake and (3) A machine learning technique to classify short intervals into sleep and awake. All methods are illustrated on a real time series in Fig 1.

**Fig 1.**
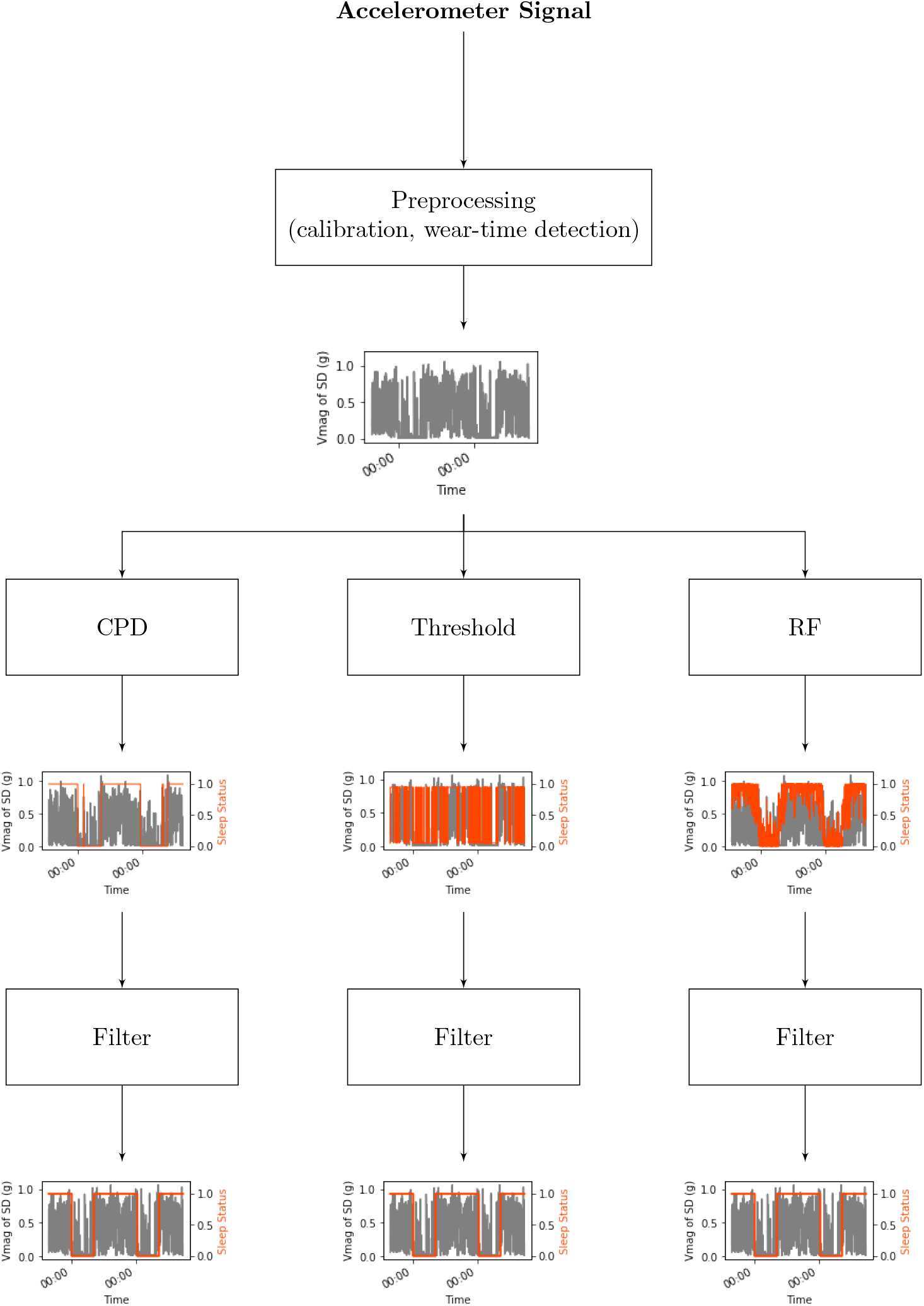
Processing steps. Visual representations automated detection of sleep-boundary times as described in this paper. (1) refers to the change-point detection based method, (2) refers to the threshold based method and (3) to the random forest based method.

### Sleep-labeling using change-point detection

Accelerometer signals during wakefulness contain more severe and frequent changes in acceleration than during sleep, resulting in differences in both the mean and variance. For the purpose of this analysis the standard deviation of acceleration along the x-, y- and z-axes were calculated over one-minute epochs and combined as the vector magnitude of standard deviations, *s_vmag*.

The change-point detection technique developed by *Killick et al* [13], implemented in R package *‘changepoint’* [14] was employed to identify changes in the mean or variance of *sjvmag.* This technique is based on maximum log-likelihood estimations (MLE), which provide a measure of the integrity/agreement within a signal. If the sum of the MLEs of the segments of a signal prior to and after a time *n* is larger than the MLE of the combined signal, *n* is marked as a change point within the signal. In order to identify multiple change points within the signal, a binary-segmentation approach was employed, whereby the dominant change point was identified first and either side of the change point was consecutively scanned for additional change points. This search stops when no more change points can be identified.

Whilst difference in acceleration between sleep and wake stages are expected to create dominant change points, there may also be changes in accelerometer patterns throughout the day or night that result in false positive change points, e.g. a subject may show predominantly active behaviours in the morning and then become more sedentary in the afternoon (Fig 2 A). In order to improve the performance of the change-point-detection based sleep classification method, a threshold (*τ_CPD_*) was included post change-point detection. Whenever the mean *s_vmag* of a section identified by the change-point-detection method fell above *τ_CPD_*, the section was classified as wakefulness, whenever the mean *sjumag* fell below *τ_CPD_*, the section was classified as sleep (Fig 2 B).

**Fig 2.**
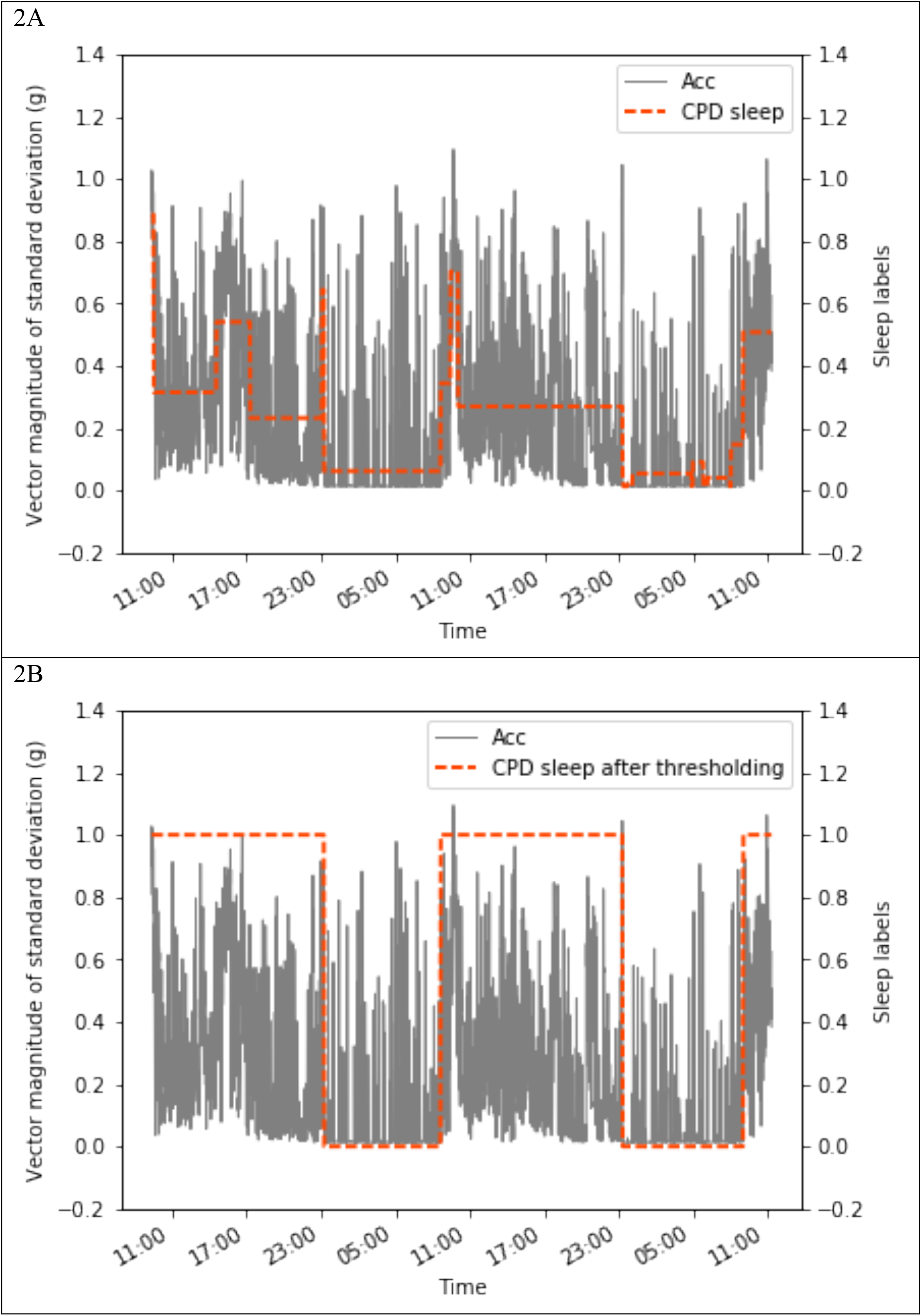
Sleep-labeling using change-point detection. A: Visualisation of false positive change point showing accelerometer signal (grey line) and mean *s_vmag* values for segments identified using change-point detection (orange dashed line). B: Accelerometer signal (grey line) and sleep (0) and wake (1) classes (orange dashed line) after thresholding.

In order to find the optimal value of *τ_CPD_*, participants were randomly assigned to a training or testing set following a 70:30 split. Thresholds between 20 mg and 150 mg were scanned in increments of 10 mg (limits were chosen based on minimum and maximum crossing points between the probability density functions of standard deviations during self-reported sleep and wakefulness). The threshold that resulted in the smallest mean absolute error (MAE) between automatically-derived and self-reported sleep-onset and wake-up times on the training set was chosen for further evaluation on the testing set.

### Sleep-labeling using thresholding

As an alternative method, a data-driven threshold was used to classify each minute of the accelerometer recording as ‘sleep’ or ‘wakefulness’. The threshold was found using probability-adjusted crossing-points of probability density functions (PDFs) during wake and sleep times (see example in Figure 3). In order to find the optimal threshold (*τ_i,j_*) for each recording in the training set, the PDFs of sleep and wake times were adjusted by their relative probabilities-of-state. Assuming an average amount of eight hours sleep a night for 24 hours worth of data, a wake-to-sleep ratio of 2:1 was used. The intersections between the adjusted wake and sleep PDFs were found and the corresponding *s_vmag_* values provided *τ_i,j_*. The PDF-based thresholds were averaged across each individual (*j*) and consecutively across all participants (*i*) in the training set in order to find the final threshold *τ_PDF_*. For the PDF-based sleep labels, all values of *s_vmag_* above *τ_PDF_* were classified as wake (1) and all others as sleep (0) (Equation 2).

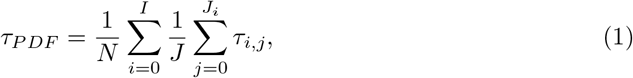

where *I* is the total number of participants in the training set and *J_i_* is the number of recordings for participant i.

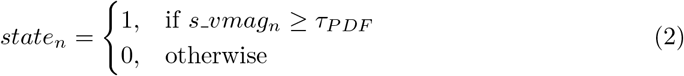

**Fig 3.**
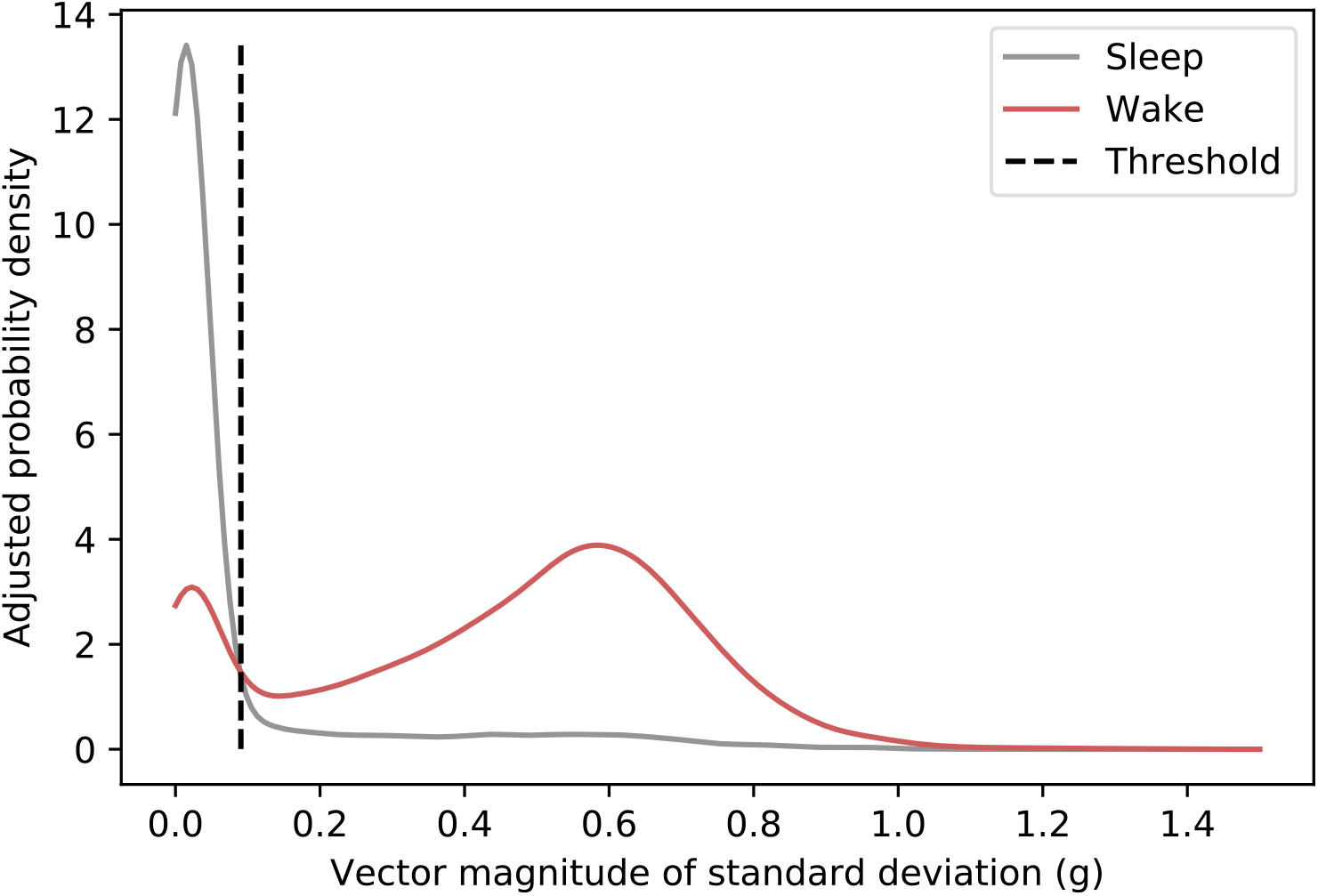
Extraction of PDF-based thresholds for sleep detection.

### Sleep-labeling using a random forest classifier

Finally, the random forest classifier developed by *Breiman* [15] and implemented in the Python package *scikit-learn* [16] was employed to classify each minute of the accelerometer recording as ‘sleep’ or ‘wakefulness’. Random forests are bagging classifiers constructed of multiple de-correlated decision trees. Each tree is itself a classifier that is fed a random subset of the dataset. During training of the random forest, the combined trees act as a form of cross-validator, reducing the classification variance due to noise within individual classifiers.

The input to the random-forest classifier was an array of 42 time- and frequency-domain features, containing single-axis features computed using the vector magnitude of acceleration *(sjvmag*) as well as cross-axial features (Table 1) [17]; extracted across one-minute epochs. In order to study the impact of age and sex on classification performance, the random-forest classifier was trained twice, once including solely accelerometer-derived features and once including accelerometer-derived features as well as age and sex information.

**Table 1.** Overview of features used for random forest classification.

| Single-axis Features | Multi-axes Features |
| --- | --- |
| Mean | Pearson correlation |
| Standard deviation | Covariance |
| Coefficient of variation |  |
| Skewness | Mean roll |
| Kurtosis | Mean pitch |
| Median | Mean yaw |
| Minimum | Standard deviation of roll |
| Maximum | Standard deviation of pitch |
| Power contained in 1-15 HZ | Standard deviation of yaw |
| Dominant frequency |  |
| Power in dominant frequency |  |
| Entropy |  |

In order to find the optimal parameters for the random-forest classifier, the training set was split into a training (*rf_train*) and a testing (*rf_test*) set (split ratio 70:30). The classifier was trained on *rf_rain* and tested on *rf_test*. Area under the curve (AUC) values for numbers of trees between 50 and 300 and minimum number of samples per leaf of 10 to 1000 were recorded for the *rf_test* set. The classification settings with the largest associated AUC were selected for further analysis.

### Post-processing using a smoothing filter

Short periods of wakefulness during sleep were sometimes classified erroneously as being the start or end of sleep. Smoothing was used to reduce this using a rolling mean-filter of length N (shown in Fig 1). N is the number of minutes to smooth over, and was learned from the training set by minimising the Mean absolute errors (MAE) between self-reported and automatically-derived sleep-onset and wake-up times post filtering. The MAE was calculated as shown in Eq 3, where *tan_i,k_* is the self-reported and *talg_i,k_* is the algorithm-derived sleep-onset or -end time, *K* is the combined number of sleep-onset and wake-up times for a specific individual and *I* the number of individuals in the training set.

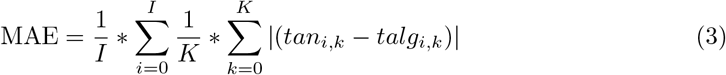

Finally, the start and end times of the *M* largest sleep periods were selected as the sleep-boundary times identified by the change-point detection method, where *M* was the number of days (defined as number of 4:00 am instances) contained within a recording.

The code developed as part of this work is available at https://github.com/activityMonitoring. (From time of publication)

### Statistical Analysis

Outcome statistics were recorded for the previously unseen test set. MAE between automatically-derived and self-reported sleep-onset and wake-up times were used as the primary outcome measure. In addition to this, kappa statistics [18] (Eq 4) between the automatically-derived and self-reported minute-by-minute labels were calculated. Kappa statistics reflect the inter-rater agreement between two sources taking into account both the observed agreement (*P_o_*, also known as accuracy) and the likelihood of them agreeing by chance (*P_e_*). Finally, Bland-Altman [19] plots were used to visualise the agreement between the automatically-derived and self-reported sleep-boundary times.

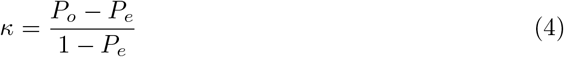

The performance of all sleep-detection algorithms was evaluated on a per person *(individual*) and a per night *(episode*) basis. For the *individual* analysis, mean values across all recording incidences and all nights were extracted.

## Results

### Study Population

Out of the 51 participants taking part in the CHA Exercise study, 46 had at least one day worth of accelerometer recordings with corresponding self-reported sleep information. A total of 972 days of accelerometer recordings were included in the analysis. The mean age of all included participants was 66.7 (standard deviation: 5.4) years. 29 (63%) participants were female, 17 (37%) male. Table 2 provides an overview of participant characteristics and self-reported data grouped by tertiles of total sleep duration (tertile boundaries: 0h-7.3h, 7.4h-8.0h, 8.1h-9.2h).

**Table 2.** Baseline characteristics and self-reported sleep information of participants by tertiles of self-reported total sleep time.

|  | Tertile 1<br>(0-7.3h) | Tertile 2<br>(7.4-8h) | Tertile 3<br>(8.1-9.2h) |
| --- | --- | --- | --- |
| n | 15 | 15 | 16 |
| Age (years) | 66.5 (5.0) | 67.9 (6.2) | 65.7 (5.2) |
| % female | 60 | 67 | 56 |
| Self-reported days | 21.9 (5.8) | 20.1 (8.0) | 20.0 (5.4) |
| Total sleep time (h) | 6.5 (1.0) | 7.7 (0.2) | 8.4 (0.4) |

### Sleep-boundary detection

The threshold to differentiate between sleep and wake episodes identified by the change-point-detection method, *τ_CPD_*, learned using the training set was 0.192g. The ideal filter length, *N*, learned using the training set was 60 minutes. The performance of the change-point detection based sleep-boundary detection on the testing set is summarised in Table 3. The method achieved a MAE of 36 min and a kappa statistic of 0.87 in the *individual* analysis. Figures 4.a-b show the Bland-Altman plots of sleep-onset and -end times.

**Fig 4.**
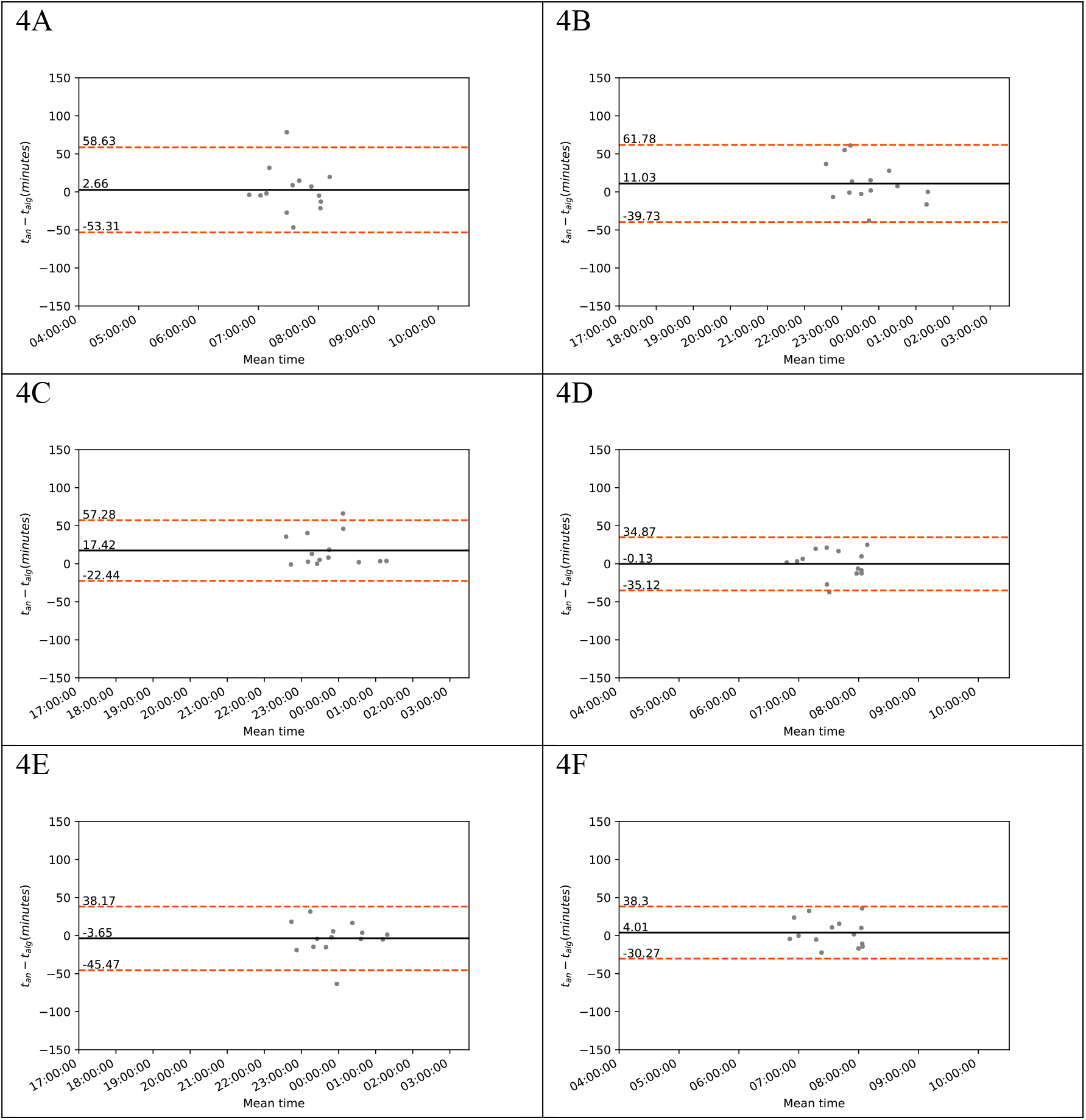
Bland-Altman plots of agreement between automatically-derived (t_alg) and self-reported (t_an) sleep-onset and wake-up times on an individual level. A: Change-point detection sleep-onset times. B: Change-point detection wake-up times. C: Thresholding sleep-onset times. D: Thresholding wake-up times. E: Random forest sleep-onset times. F: Random forest wake-up times.

**Table 3.** Performance summary: Mean absolute error (MAE) and kappa statistic between automatically-derived and self-reported sleep-onset and wake-up times.

|  | Change-point detection<br>(minutes) | Thresholding<br>(minutes) | Random forest<br>(minutes) |
| --- | --- | --- | --- |
| **Individual (n=14)** |  |  |  |
| MAE combined (SD) | 35.9 (14.8) | 34.4 (9.9) | 33.3 (11.3) |
| MAE wake (SD) | 20.3 (20.3) | 14.9 (9.9) | 14.6 (10.5) |
| MAE sleep (SD) | 20.3 (19.5) | 17.6 (20.2) | 14.7 (15.9) |
| MAE TST (SD) | 31.7 (32.0) | 25.7 (18.0) | 19.3 (14.7) |
| Kappa statistic | 0.87 | 0.89 | 0.89 |
| **Episode (n=264)** |  |  |  |
| MAE combined (SD) | 31.8 (54.8) | 32.8 (38.0) | 31.1 (41.5) |
| MAE wake (SD) | 26.3 (43.4) | 30.2 (33.3) | 27.8 (34.3) |
| MAE sleep (SD) | 37.4 (47.4) | 35.4 (42.0) | 34.5 (47.3) |
| MAE TST (SD) | 36.4 (45.9) | 39.2 (37.8) | 36.7 (40.6) |
| Kappa statistic | 0.89 | 0.89 | 0.90 |

The optimal data-driven threshold (*τ_PDF_*) to classify each minute of the accelerometer recording as ‘sleep’ or ‘wakefulness’ was found to be 0.074g. The corresponding filter length N for the thresholding method was found to be 70 minutes. Using these settings the thresholding method achieved a MAE of 34 min and a kappa statistic of 0.89. Figures 4.c-d show the Bland-Altman plots of sleep-onset and -end times.

**Fig 5.**
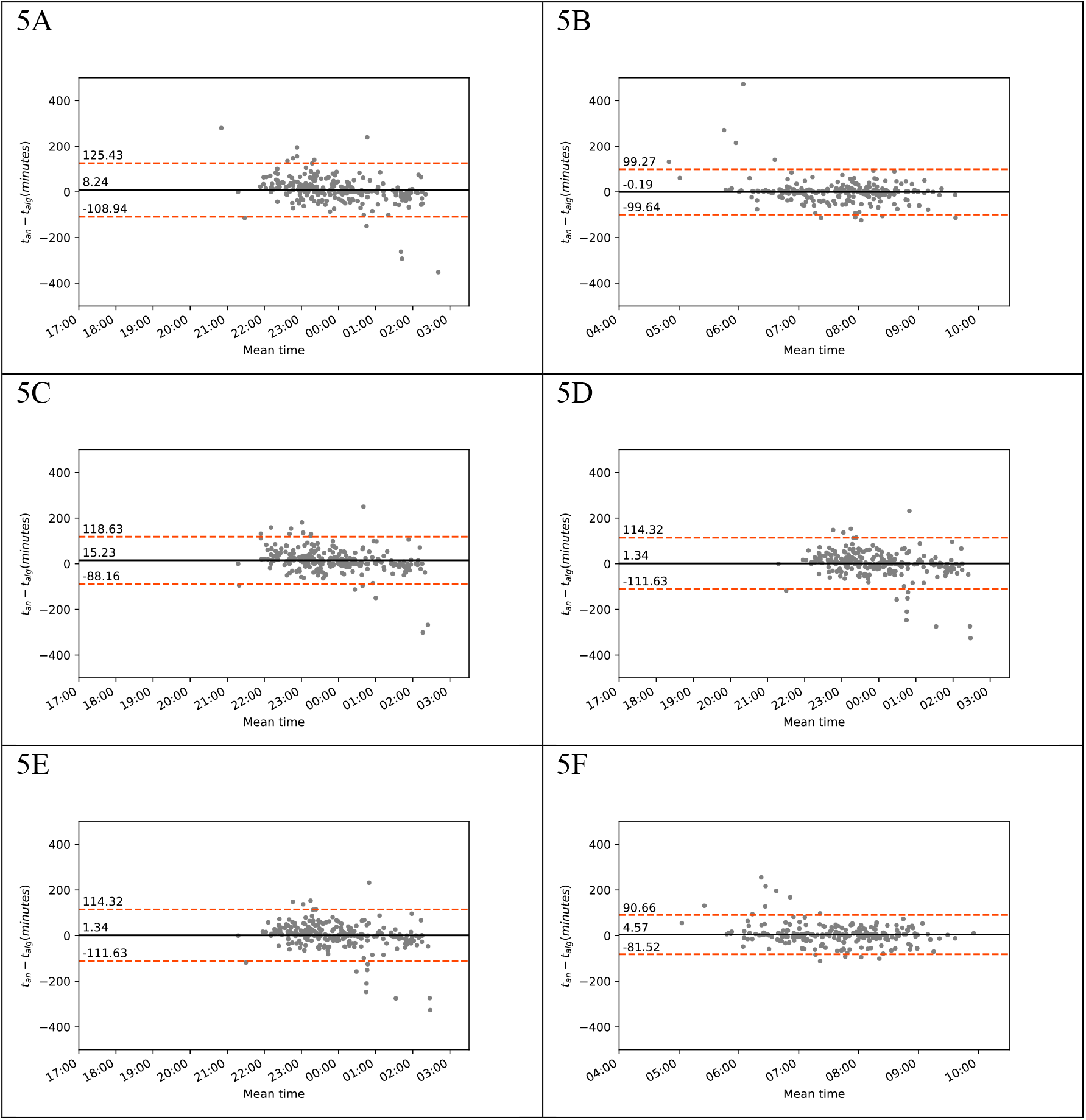
Bland-Altman plots of agreement between automatically-derived (t_alg) and self-reported (t_an) sleep-onset and wake-up times on an episode level. A: Change-point detection sleep-onset times. B: Change-point detection wake-up times. C: Thresholding sleep-onset times. D: Thresholding wake-up times. E: Random forest sleep-onset times. F: Random forest wake-up times.

The feature importance learned by the random forest classifier excluding age and sex information is shown in Figure S1 Fig in the Appendix. Median, 75th percentile and 25th percentile were identified as the most important features. The choice of maximum number of trees and minimum numbers of leafs only minimally affected the results; the combination of 10 trees and a minimum of 10 leafs resulted in an area under the curve (AUC) value for minute-by-minute sleep classification of 0.95. The ideal filter length *(N*) was found to be 50 minutes, achieving a MAE of 33 minutes and a kappa statistic of 0.89 in the *individual* analysis (see Table 3).

Including age and sex information as features in the classification did not improve the method’s performance, resulting in a MAE of 32.1 (40.1) and a kappa statistic of 0.89. The feature importance learned by the classifier including age and sex information is shown in Figure S2 Fig in the Appendix.

The Bland-Altman plots for both the change-point detection and random forest based classification are shown in Figure 4. They show on average good agreement between self-reported and automatically-derived sleep-boundary times. There are however outliers with more than 100 minutes difference between self-reported and automatically-derived sleep boundary times. On closer inspection, these outliers include potential errors in self-reported sleep-boundary times (mislabeling) as well as miss-classifications of sleep and wake labels. Miss-classifications occur primarily where low-activity wake times and sleep times are close in proximity and when sleep-interruptions occur in close proximity to sleep-boundary times.

## Discussion

To our knowledge, this is the first study attempting a data-driven approach to automated sleep-boundary detection. Previous studies have used self-reported sleep times [6] or restricted their analysis to night-time assessments [20]. As a result, current accelerometer studies are reliant on the collection of self-reported sleep-boundary information. Where this information is missing, analysis of sleep and wake behaviour of participants is severely limited. The proposed sleep-boundary detection methods allow for the identification of sleep onset and end times in the absence of self-reported data and open the door to large-scale fully-automated accelerometer analysis.

The methods were trained using self-reported sleep and wake information as their ground truth. Self-reported sleep times have been shown to be less accurate than objectively collected sleep-onset and end-times [5] and may be seen to provide a weaker ground truth than polysomnography assessments. Collecting free-living rather than laboratory data does however provide a more realistic setting for sleep analysis, where recordings may be affected by periods of rest prior to sleep and after wake-up that may otherwise be missed. In addition to this, allowing participants to stay in their natural sleeping environment reduces the impact of data collection on their sleeping behaviour [21].

Previous research by *Lauderdale et al* has shown that self-reported sleep information can be affected by mis-reporting [5]. Whilst some differences may be explained by errors in self-reporting [5], others are likely to be caused by miss-classifications. Such miss-classifications have in the past been reported in cases where sleep episodes were followed by restful wake periods [22].

The algorithms described as part of this work have been developed using raw accelerometer data as opposed to proprietary count-values. Nevertheless, device- (e.g. on-board filtering) and protocol-specific (e.g. wearing location) characteristics need to be considered when applying the developed algorithms to new datasets.

## Conclusion

We developed three automated sleep-boundary detection methods for the analysis of accelerometer data that will not need self-reported sleep-diary information to label new accelerometer data. The change-point detection, thresholding and random forest methods achieved MAE of 36 min, 34 min and 33 minutes, and kappa statistics of 0.87, 0. 89 and 0.89, respectively. A kappa statistic of 0.7 and above is traditionally thought of as a substantial level of agreement [18]. Whilst the random forest method achieved the lowest MAE, the thresholding method performed only marginally worse and requires only a single input variable. We therefore recommend this approach for automated sleep-boundary detection. This work can facilitate the inclusion of polysomnography-validated measures of sleep status in population scale studies.

## Supporting information

**S1 + S2 Fig.**
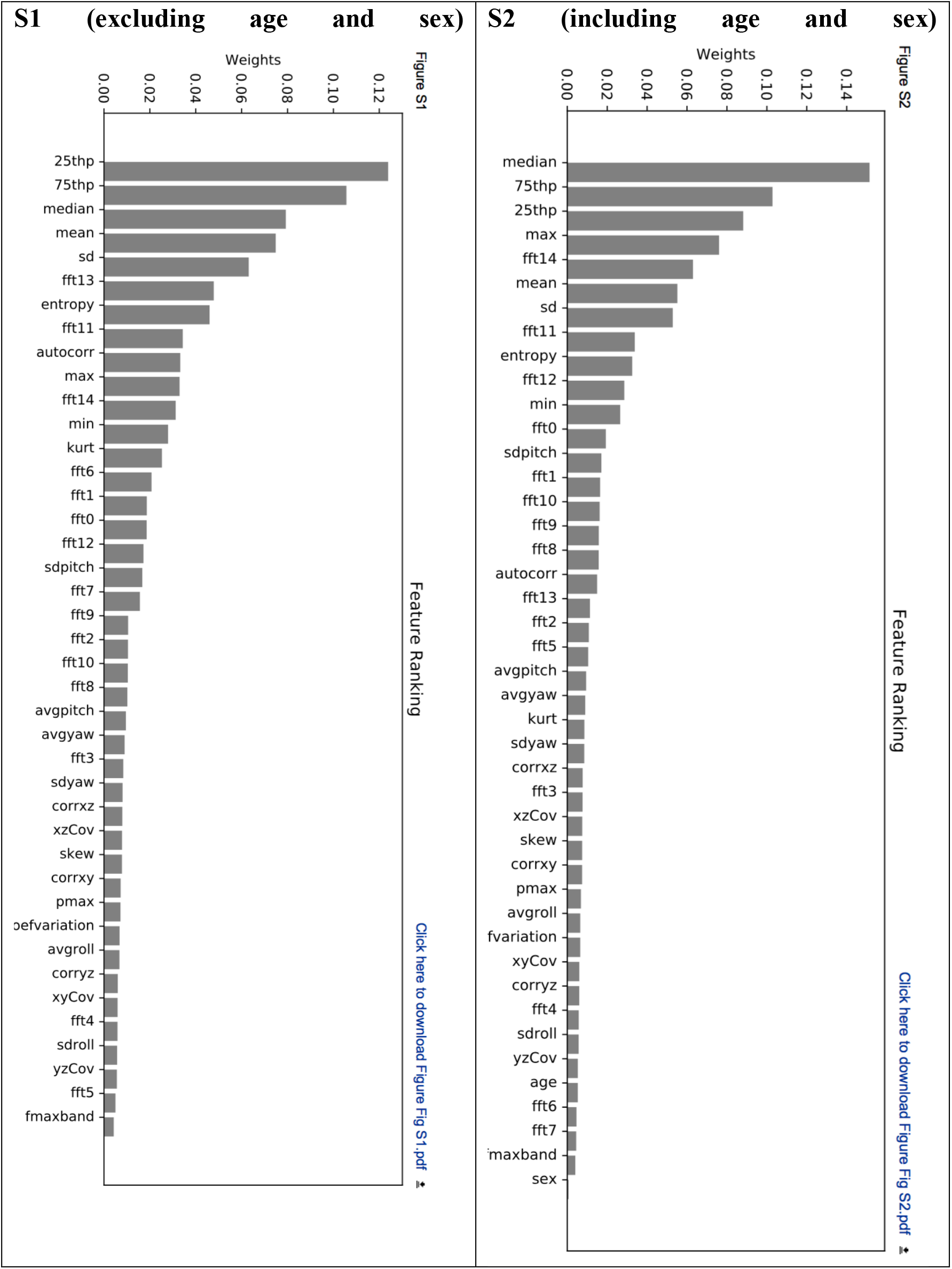
Feature ranking as learned by random forest classifier.

## Acknowledgments

We would like to thank all participants for agreeing to volunteer in this research, and Jill Betts for organising this data collection effort. This analysis was supported by the British Heart Foundation Centre of Research Excellence at Oxford [grant number RE/13/1/30181 to AD], the National Institute for Health Research (NIHR), Oxford Biomedical Research Centre (BRC) based at Oxford University Hospitals NHS Trust and University of Oxford, the NIHR Oxford Health BRC and the Wellcome Trust. We would also like to acknowledge the use of the University of Oxford Advanced Research Computing (ARC) facility in carrying out this work (http://dx.doi.org/10.5281/zenodo.22558). JOD acknowledges the support of the RCUK Digital Economy Programme with grant number EP/G036861 (Oxford Center for Doctoral Training in Healthcare Innovation). No funding bodies had any role in the analysis, decision to publish, or preparation of the manuscript.

